# MS/MS fragmentation pattern analysis confirms the production of the new esterified bile acids by the human gut microbiota

**DOI:** 10.1101/2023.11.07.564921

**Authors:** Carlos J. Garcia, Rocio García-Villalba, David Beltrán, Maria D. Frutos-Lisón, Francisco A. Tomás-Barberán

**Affiliations:** Quality, Safety and Bioactivity of Plant-Derived Foods, Centro de Edafología y Biología Aplicada del Segura-Consejo Superior de Investigaciones Científicas (CEBAS-CSIC), 30100 Murcia, Spain

**Author notes:** Corresponding author: (ext. 445454).

**Keywords:** Bile acids, MCBAs, Gut microbiota, MS/MS spectra

## Abstract

The pathophysiology of bile acids (BAs) has been widely studied. The BAs are signaling molecules that affect lipids and glucose homeostasis via activation of BAs FRX and TGR5 receptors in the liver and peripheral tissues. The ratio of conjugated/unconjugated BAs seems relevant to quantify these interactions and, therefore, the impact on the metabolism. The gut microbiota plays a key role because they convert the primary BAs into the secondary BAs, hydrolyzes the hepatically conjugated BAs and re-conjugates BAs with amino acids. New re-conjugated BAs with amino acids (MCBAs) in the form of amides or esters have been recently suggested, but it was not possible to confirm them. This study evaluates the production of MCBAs by human gut microbiota in in vitro colonic fermentations and designs a bioanalytical method to discriminate between amides and esters. Amides and seven new esters of re-conjugated BAs composed of lithocholic acid conjugated with leucine, valine and aminobutyric acid were identified and confirmed by MS/MS after incubation with chenodeoxycholic acid and lithocholic acid. There were no specific fragments in negative polarity to discriminate between amides and esters. However, in positive polarity the amides showed a characteristic MS/MS fragment consisting of the loss of water from the released amino acid, while the esters showed the loss of water plus carbon monoxide. This study confirmed for the first time the presence of esterified MCBAs, in addition to amides, and characterized the specific MS/MS fragmentation patterns to identify and discriminate them. These results show for the first time the existence of re-conjugated BAs by ester bond and the capability to produce them by the gut microbes. This bioanalytical method will allow including these new MCBAs in the BAs analysis.

## Introduction

Bile acids (BAs) are endogenous steroids widely studied because they are closely associated to human health.^**1**^ BAs are terpenoids biosynthesized in the hepatocytes from cholesterol after several enzymatic reactions grouped in two different pathways.^**2**^ These reactions lead to primary BAs, which include cholic acid (CA) and chenodeoxycholic acid (CDCA). The primary BAs are then conjugated with glycine and taurine by bile acid-CoA amino acid N-acyltransferase (BAAT) and stored in the gall bladder until they are released into the intestine as a response to food intake.^**3**^ The role of BAs is key in lipid metabolism and because of their amphiphilic nature, they have an essential function in fat absorption. Further, BAs are also signaling molecules and metabolic regulators that activate the nuclear receptor and G protein-coupled receptor signaling to regulate hepatic lipids, glucose, and maintain metabolic homeostasis.^**4**^ Much effort has been dedicated to understanding the complex pathophysiology of the BAs, but they still need to be completely unravelled. The interaction between the gut microbiota and the BAs is the crucial point that makes this understanding difficult. Once the conjugated primary BAs are released, through the bile duct, to the intestine they interact with the gut microbiota converting them into secondary bile acids.^**5**^ The secondary BAs include the dehydroxylated, oxidized (oxo-BAs) and epimerized BAs.^**6**^ This modifies the final pool of the BAs giving a large family variability and alters their final biological effect. The gut microbes usually perform deconjugation, dehydroxylation, dehydrogenation, and epimerization of the BAs but it has been recently discovered the occurrence of a new transformation, the re-conjugation with amino acids leading to metabolites known as “microbially conjugated bile acids” (MCBAs).^**7**^ This greatly expands the variety of the secondary BAs family and its implications.^**8**^ Initially, the term MCBAs only referred to amides conjugated with amino acids in the C24 acyl site similar to those conjugated with glycine and taurine in the hepatocyte. More recently, we suggested a new possible BAs re-conjugation by an ester bond on the hydroxyl groups of the sterols backbone.^**9**^ In this study, five isomers of lithocholic (LCA) or isolithocholic (iLCA) conjugates with valine were observed, however only four conjugations with amide bond would be theoretically feasible considering the valine enantiomers, suggesting the existence of a new type of conjugate. However, both amides and esters showed apparently the same MS/MS fragmentation spectra releasing the amino acids, and therefore it was not possible to differentiate between them and confirm their presence. In the same way either the mechanisms by which they are produced and the biological effects of these new re-conjugated BAs are still unknown. The previous study suggests the occurrence of MCBAs as a bacterial mechanism to reduce the toxicity of the unconjugated BAs as consequence of a high incidence of the Bile salt hydrolase (BSH). The BSH activity is carried out by the anaerobic intestinal bacteria of the genera Bacteroides, Clostridium, Lactobacillus, and Bifidobacterium.^**10**^ BAs in general and conjugated BAs have been correlated with relevant metabolic disorders such as dyslipidemia, hypercholesterolemia and dysglycemia.^**11,12,13**^ This correlation exists because the balance of the conjugated/unconjugated BAs in the ileum may alter the regular interaction with intestinal receptor GLP-1 and stimulate the insulin secretion, as well as inhibit the intestinal FXR-FGF15 signaling pathway reducing hepatic cholesterol and decreasing lipogenesis.^**14**^ BAs are signaling molecules as it has been documented in many studies.^**15,16**^ It has been demonstrated that primary BAs and secondary BAs are signaling molecules, acting on several receptors including the G protein-coupled bile acid receptor 1 (GPBAR1 or Takeda G-protein receptor 5) and the Farnesoid-X-Receptor (FXR). Because of this, the BAs have been associated with metabolic disorders related to cholesterol, lipids and glucose metabolism, bowel disease or cancer .^**17,18,19,20,21,22,23,24**^ These studies point to the gut microbes as a key to go one step further in knowing the pathophysiology of the secondary BAs. The evidence highlights the need to enlarge the knowledge about this large family of MCBAs and to develop methods to identify the new MCBAs produced by esterification and to discriminate them from the amides in order to quantify the total amount and type of the conjugated BAs.

This study aims to optimize a bioanalytical method by LC-MS to analyze the new re-conjugated BAs in fecal samples and in in vitro fecal incubations and investigate thoroughly the MS/MS fragmentation patterns of MCBAs to confirm their structures and validate their different chemical polarities.

## Materials and Methods

### Chemicals

Acetonitrile and water 0.1% (v/v) formic acid were purchased from J.T. Baker (Deventer, The Netherlands), and formic acid was obtained from Panreac (Barcelona, Spain). Authentic standards of 3,7-Dihydroxy-5-cholan-24-oic Acid (chenodeoxycholic acid) and 3-Hydroxy-11-oxo-5-cholan-24-oic Acid (3-oxo-chenodeoxycholic acid) were purchased from Avanti Polar Lipids (Alabaster, AL, USA). The N-(3α,7α,12α-trihydroxy-5β-cholan-24-oyl)-glycine (glycocholic acid), N-(3α,7α,12α-trihydroxy-5β-cholan-24-oyl)-taurine (taurocholic), 3α,7α,12α-trihydroxy-5β-cholan-24-oic acid (cholic acid), 3α,7β,12α-Trihydroxy-5β-cholan-24-oic acid (ursocholic), 3α,6β-Dihydroxy-5β-cholan-24-oic acid (murideoxycholic acid), 3α,6α-Dihydroxy-5β-cholan-24-oic acid (hyodeoxycholic acid), 3α,7β-Dihydroxy-5β-cholan-24-oic acid (ursodeoxycholic acid), 3α,12α-Dihydroxy-5β-cholan-24-oic acid (deoxycholic acid), 3α-Hydroxy-5β-cholan-24-oic Acid (lithocholic) and 3β-Hydroxy-5β-cholan-24-oic Acid (isolithocholic) were purchased from Cayman Chemical (Ann Arbor, MI, USA).

### Collection of human fecal samples

Three healthy donors of stool samples, two male (age 45, 35) and a female (age 32), were recruited at the Centro de Edafología y Biología Aplicada del Segura (CEBAS-CSIC, Murcia, Spain) and all gave written informed consent. The sample evaluation of three donors is adequate to describe the presence and confirmation by MS/MS fragmentation metabolomics of re-conjugated BAs.

### In vitro incubation of fecal samples

Preparation of fecal suspensions and subsequent fermentation experiments were performed similarly to previous studies with brief modification.^**9**^ The fermentation procedure was performed under anoxic conditions in an anaerobic chamber (Concept 400, Baker Ruskinn Technologies, Ltd., Bridgend, South Wales, UK) with an atmosphere consisting of N2/H2/CO2 (85:5:10) at 37 °C. Aliquots of stool samples (10 g) were diluted 1/10 w/v in Nutrients Broth supplemented with 0.05% l-cysteine hydrochloride and homogenized by a stomacher in filter bags. Different types of samples were used: i) BAs + fecal samples incubated; ii) fecal samples incubated (control 1); iii) BAs + fecal samples non-incubated; and iv) fecal samples non-incubated (control 2). Aliquots of fecal suspensions (50 µL) were inoculated into 5 mL of fermentation medium anaerobe Wilkins Chaldean containing 50 µM of chenodeoxycholic acid (CDCA) or lithocholic acid (LCA). The chenodeoxycholic acid was selected because is a primary BA with medium hydrophobicity and is used by the gut bacteria to produce the secondary BAs. In previous studies, bacteria converted CDCA into LCA, which is more hydrophobic and cytotoxic, and then the gut microbes reduce its hydrophobicity by re-conjugations it with amino acids. The LCA was used to confirm if the MCBAs (esters and amides) are produced from the LCA or from its isomers or epimers. Three replicate cultures were prepared in parallel from each fecal suspension. Samples were collected at 0, 24,48,72,96 and after 120 hours (five days) of incubation at 37 °C. The duration of the fecal incubation was set in order to ensure conversion procedures by the gut microbiota. Usually after 24–48 the bacteria are already in a stationary phase and it is in this phase when the secondary metabolism, which acts in this conversion, commonly occurs .^**25**^

### BAs extraction for metabolomics analysis

Prior to the UPLC-ESI-QTOF-MS, the samples were extracted according to previous studies with modification due to liquid samples after 5 days of in vitro incubation were used instead of fresh fecal samples as usual **(Fig. 1)**.^**26**^ 5mL of incubation solution were vortexed and centrifuged at 4500 rpm for 15 min at 4 °C. Then an SPE extraction was performed using a cartridge Hypersep Sep 500mg/2.8mL C18 by using manifolds. The cartridge was then washed with water (5 mL) and bile acids were eluted with ethanol (5 mL).

**Figure 1.**
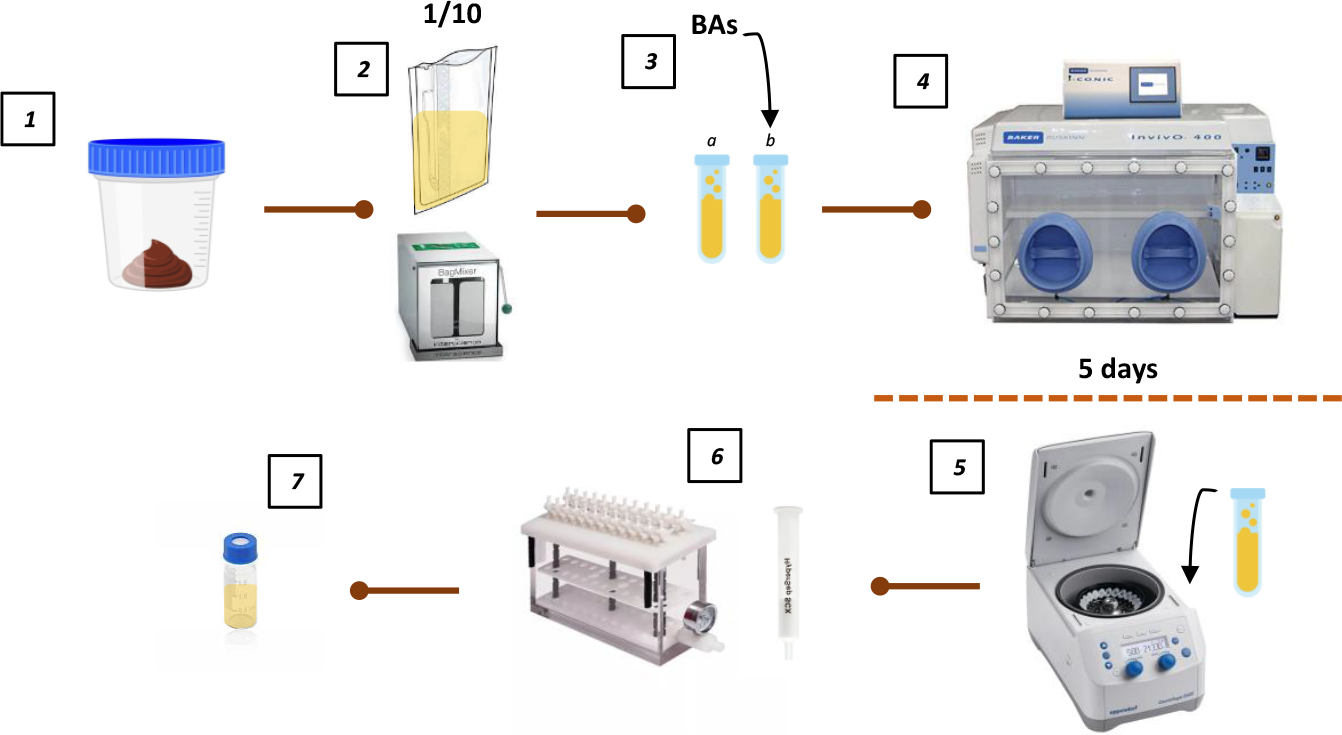
BAs analysis workflow. 1) Fresh fecal samples collected by each volunteers; 2) Dilution 1/10 of the sample in the medium and mixture by stomacher during 3 minutes; 3) Incubation tubes including: medium + total fecal bacteria (a), medium + total fecal bacteria + BAs (b); 4) Tubes were incubated during 5 days at anaerobic condition; 5) After each sample point samples were centrifuged and supernatant collected; 6) SPE extraction were performed for BAs concertation; 7) The volume eluted were dry by speed vacuum and re-dissolved in 100 µl of MeOH prior UPLC-QTOF-MS analysis.

### UPLC-QTOF-MS analysis

The metabolomics analysis was performed on a U-HPLC (Infinity 1290; Agilent) coupled to a high-resolution mass spectrometer with a quadrupole time-of-flight mass analyzer (6550 iFunnel Q-TOF LC/MS; Agilent) with an Agilent Jet Stream (AJS) electrospray (ESI) source. The mass analyzer was operated in negative mode under the following conditions: gas temperature 150° C, drying gas 14 L/min, nebulizer pressure 40 psig, sheath gas temperature 350° C, sheath gas flow 11 L/min, capillary voltage 3500 V, fragmentor voltage 120 V, and octapole radiofrequency voltage 750 V. Data were acquired over the m/z range of 50–1700 at the rate of 3 spectra/s. The m/z range was autocorrected on reference masses 112.9855 and 1033.9881.

The MS/MS target product ion spectra were acquired at m/z 100–1100 using a retention time window of 1 min, a range of 5-60 eV of collision energy an acquisition rate of 1 spectra/s. The chromatographic analysis was performed with a reversed-phase C18 column (Poroshell 120, 3 x 100 mm, 2.7 µm pore size) at 30° C, using water + 0.1% formic acid (Phase A) and acetonitrile + 0.1% formic acid (Phase B) as mobile phases with a flow rate of 0.4 mL/min. The gradient started with 50% B, increased in 4 min to 90% B, in 3 min to 99% B, held for 3 min and decreased to the initial conditions during 3 min. The injection volume for all samples was 3 µL. Raw data acquired was processed by MS-DIAL 5.1.2 (prime.psc.riken.jp/compms) through the application of a featured extraction based on the in-house database built for bile acids. After data processing, the bile acids identified were analyzed by Mass Hunter Qualitative 10.0 qualitative (Version B.10.0, Agilent software metabolomics, Agilent Technologies, Waldbronn, Germany).

### Results and discussion

The bioanalytical method proposed in this study was designed to distinguish the MCBAs, in particular the new esters from the regular amides similar to those conjugated hepatically with glycine and taurine. To this goal, both the extraction, chromatography and MS parameters were especially set to increase the equipment response to BAs. This study evaluated the bacterial metabolism of the CDCA and LCA during 120 hours to produce the secondary BAs by the classical mechanisms and the new re-conjugated BAs. As previously described, after the incubation CDCA was majorly converted to LCA and its epimer (iLCA), and isoursodeoxycholic acid (isoUDCA) and ursodeoxycholic acid (UDCA) accumulating them after 5 days of incubations.^9^ The fermentation results showed a global trend to accumulate the monohydroxylated BAs and epimers of CDCA via mostly 7α hydroxyl position metabolism instead of 3α **(Fig. 2)**. The results were able to show the pathway intermediates because the study included sampling points every 24 h. The CDCA, LCA, iLCA (not in all volunteers) and 7α-Hydroxy-3-oxo-5β-cholan-24-oic Acid (3-oxo-chenodeoxycholic Acid or 3-oxo-CDCA) were detected in fecal samples at time 0. LCA and iLCA increased after incubation, however the 3-oxo-CDCA showed a decrease during the fermentation of CDCA. Therefore, its occurrence from CDCA was not relevant and its derivatives (Grey box) were mainly produced from the fecally excreted 3-oxo-CDCA.The identified secondary BAs derived from the bacterial metabolism of CDCA were: i) LCA (dehydroxylated BAs); ii) 7-oxoLCA acid, 3-oxoCDCA acid, 3-oxo-5β-cholan-24-oic-Acid, 7-oxo-5β-cholan-24-oic-Acid (oxo-BAs) ; iii) iDCA, UDCA, iUDCA and iLCA (epimers BAs) **(Table 1)**. The 7-oxo-5β-cholan-24-oic-Acid was identified approximately fifteen times less intense than 3-oxo-5β-cholan-24-oic-Acid and the 7-oxo-5β-cholan-24-oic-Acid appeared after 48h of incubation and decrease. The precursor of 7-oxo-5β-cholan-24-oic-Acid, the monohydroxylated 7α-5β-cholan-24-oic-Acid, and its epimer were not detected. These results suggested a fast conversion of 7α-5β-cholan-24-oic-Acid to 7-oxo-5β-cholan-24-oic-Acid and the re-conjugation of the epimer. However, this possibility was rejected because a non-significant decreased of the 7-oxo-5β-cholan-24-oic-Acid was observed. In the case of LCA incubation, no secondary BAs were identified derived from the LCA bacterial metabolism and it was observed a decreasing of LCA along sampling points.

**Figure 2.**
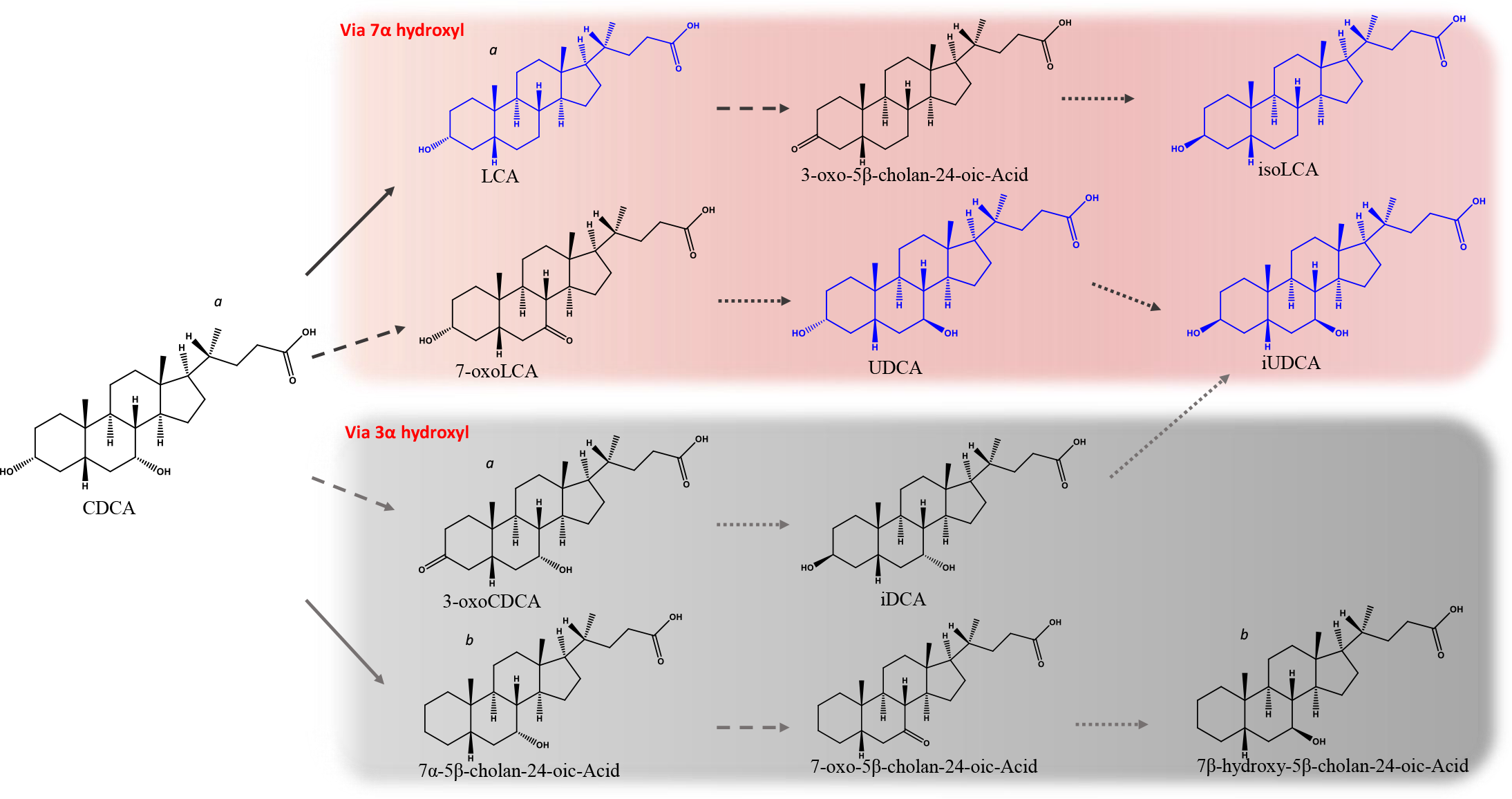
Global trend of CDCA bacterial metabolism. Red box: CDCA bacterial metabolism of 7α hydroxyl; Grey box: CDCA bacterial metabolism of 3α hydroxyl. →: Bacterial enzymatic activity of 3/7α dehydroxylase;⁃⁃⁃: Bacterial enzymatic activity of 3/7α hydroxysteroid dehydrogenase (7/3 α/β-HSDH); •••: Bacterial enzymatic activity of 3/7 α/ β hydroxysteroid dehydrogenase (7/3 α/β-HSDH). Blue BAs molecules: Bile acids accumulated produced after the incubation; Black BAs molecules: Bile acids identified during the incubation. (a) Identified in fecal samples before incubation; (b) Not identified

**Table 1.** Primary and secondary bile acids identified and confirmed by authentic standard and MS/MS fragmentation patterns.

| Compound name | Formula | m/z | ppm | Rt | MS/MS fragments (neg polarity) | Collision E |
| --- | --- | --- | --- | --- | --- | --- |
| Chenodeoxycholic acid (CDCA)<sup>a</sup> | C<sub>24</sub>H<sub>40</sub>O<sub>4</sub> | 391.2859 | 1.32 | 3.4 | 391.2862;373.2750 | 40 |
| Lithocholic acid (LCA)<sup>a</sup> | C<sub>24</sub>H<sub>40</sub>O<sub>3</sub> | 375.2922 | 4.6 | 5.18 | 375.2939;357.2838;355.2669 | 50 |
| 7-oxo-lithocholic acid (7-oxoLCA) | C<sub>24</sub>H<sub>38</sub>O<sub>4</sub> | 389.2712 | 3.76 | 2.8 | 389.2722;345.2808;343.2657 | 40 |
| 3-oxo-chenodeoxycholic acid (3-oxoCDCA)<sup>a</sup> | C<sub>24</sub>H<sub>38</sub>O<sub>4</sub> | 389.2711 | 3.5 | 3.68 | 389.2708;345.2813;343.2664;<br>371.2643 | 40 |
| 3-oxo-5β-cholan-24-oic-Acid | C<sub>24</sub>H<sub>38</sub>O<sub>3</sub> | 373.2751 | 0.75 | 5.35 | 373.2737;355.2634 | 30 |
| 7-oxo-5β-cholan-24-oic-Acid | C<sub>24</sub>H<sub>40</sub>O<sub>4</sub> | 373.2759 | 2.89 | 4.69 | 373.2740;355.2638 | 30 |
| Ursochenodeoxycholic acid (iUDCA) | C<sub>24</sub>H<sub>40</sub>O<sub>4</sub> | 391.2851 | -0.72 | 2.37 | 391.2859; 373.2780 | 40 |
| Isochenodeoxycholic acid (iDCA) | C<sub>24</sub>H<sub>40</sub>O<sub>4</sub> | 391.2850 | -0.98 | 2.73 | 391.2858;373.2755 | 40 |
| Isolithocholic acid (iLCA) | C<sub>24</sub>H<sub>40</sub>O<sub>3</sub> | 375.2916 | 3.01 | 4.5 | 375.2920; 357.2808;355.2650 | 50 |
| Isoursochenodeoxycholic acid (iUDCA) | C<sub>24</sub>H<sub>40</sub>O<sub>4</sub> | 391.2849 | -1.23 | 2.2 | 391.2855;373.2784 | 40 |

<sup>a</sup>Identified in fecal samples before incubation <sup>b</sup>Not identified

The bacterial re-conjugation of CDCA, LCA and their secondary BAs were studied. MCBAs were identified in the fecal samples and samples incubated. The MCBAs identified were secondary BAs re-conjugated with the amino acids leucine, valine and aminobutyric acid by ester and amide bonds. The LCA and isoLCA were the BAs backbone identified as part of MCBAs even after the incubation with CDCA. The non-polar amino acids valine and leucine and the non-protein amino acid aminobutyric acid were identified as the main amino acids re-conjugated BA derived from the LCA backbone **(Fig. 3)**. The LCA incubations were able to certify the MCBAs derived specifically from LCA and not from isoLCA. Three MCBAs leucine-derived were identified after incubation with CDCA and LCA; three MCBAs valine-derived were identified after the incubation with LCA instead of five with CDCA; and two MCBAs aminobutyric acid-derived were identified after the incubation with LCA instead of five with CDCA **(Table 2)**. This made it possible to confirm the occurrence of these MCBAs derived from LCA and allowed the discrimination of the MCBAs derived from LCA and isoLCA. Several isomers of valine, leucine, corresponding to the enantiomers of L and D were found. It was possible to discriminate between valine isomers because the positions of the amino group confers polarity differences previously reported being higher polar of D than L enantiomer.^**27**^ In case of aminobutyric acid re-conjugated, many options are feasible because of the position of the amino group. This position determines the polarity of the molecule being the polarity order α >β >γ.28 Additionally to these polarity features, the amides presents a theoretically more polar behaviour than esters. This made it possible to distinguish between two different isomers of MCBAs derived from the same BA backbone.

**Figure 3.**
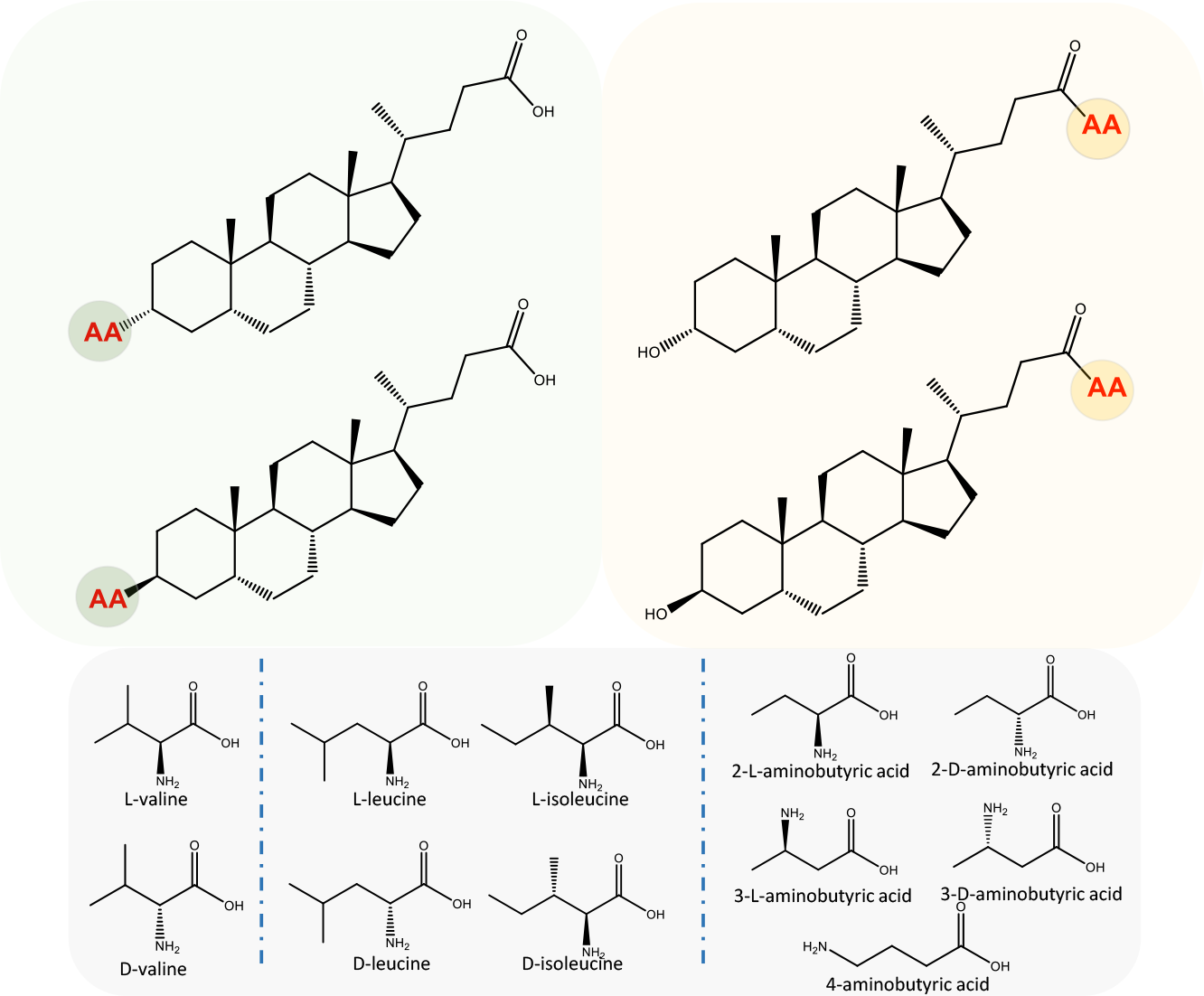
Possible combinations of MCBAs with valine, leucine and aminobutyric acid derived from the lithocholic acid backbone. Esterified MCBAs (green box); amidated MCBAs (orange box); amino acids isomers of valine, leucine and aminobutyric acid (grey box).

**Table 2.**
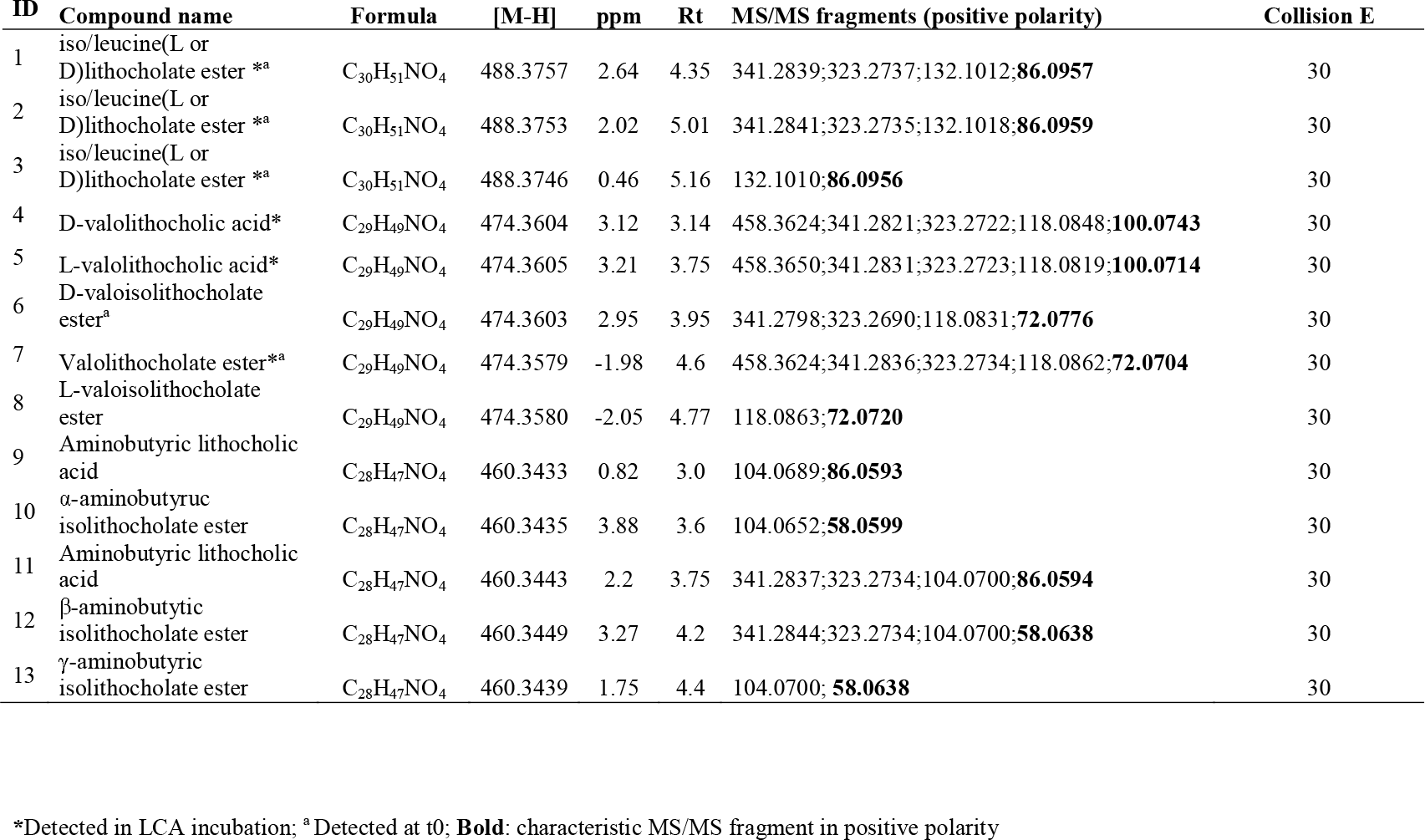
MCBAs identified and confirmed by MS/MS fragmentation patterns.

| ID | Compound name | Formula | [M-H] | ppm | Rt | MS/MS fragments (positive polarity) | Collision E |
| --- | --- | --- | --- | --- | --- | --- | --- |
| 1 | iso/leucine(L or D)lithocholate ester <sup>*a</sup> | C<sub>30</sub>H<sub>51</sub>NO<sub>4</sub> | 488.3757 | 2.64 | 4.35 | 341.2839;323.2737;132.1012;**86.0957** | 30 |
| 2 | iso/leucine(L or D)lithocholate ester <sup>*a</sup> | C<sub>30</sub>H<sub>51</sub>NO<sub>4</sub> | 488.3753 | 2.02 | 5.01 | 341.2841;323.2735;132.1018;**86.0959** | 30 |
| 3 | iso/leucine(L or D)lithocholate ester <sup>*a</sup> | C<sub>30</sub>H<sub>51</sub>NO<sub>4</sub> | 488.3746 | 0.46 | 5.16 | 132.1010;**86.0956** | 30 |
| 4 | D-valolithocholic acid* | C<sub>29</sub>H<sub>49</sub>NO<sub>4</sub> | 474.3604 | 3.12 | 3.14 | 458.3624;341.2821;323.2722;118.0848;**100.0743** | 30 |
| 5 | L-valolithocholic acid* | C<sub>29</sub>H<sub>49</sub>NO<sub>4</sub> | 474.3605 | 3.21 | 3.75 | 458.3650;341.2831;323.2723;118.0819;**100.0714** | 30 |
| 6 | D-valoisolithocholate ester<sup>a</sup> | C<sub>29</sub>H<sub>49</sub>NO<sub>4</sub> | 474.3603 | 2.95 | 3.95 | 341.2798;323.2690;118.0831;**72.0776** | 30 |
| 7 | Valolithocholate ester<sup>*a</sup> | C<sub>29</sub>H<sub>49</sub>NO<sub>4</sub> | 474.3579 | -1.98 | 4.6 | 458.3624;341.2836;323.2734;118.0862;**72.0704** | 30 |
| 8 | L-valoisolithocholate ester | C<sub>29</sub>H<sub>49</sub>NO<sub>4</sub> | 474.3580 | -2.05 | 4.77 | 118.0863;**72.0720** | 30 |
| 9 | Aminobutyric lithocholic acid | C<sub>28</sub>H<sub>47</sub>NO<sub>4</sub> | 460.3433 | 0.82 | 3.0 | 104.0689;**86.0593** | 30 |
| 10 | α-aminobutyric isolithocholate ester | C<sub>28</sub>H<sub>47</sub>NO<sub>4</sub> | 460.3435 | 3.88 | 3.6 | 104.0652;**58.0599** | 30 |
| 11 | Aminobutyric lithocholic acid | C<sub>28</sub>H<sub>47</sub>NO<sub>4</sub> | 460.3443 | 2.2 | 3.75 | 341.2837;323.2734;104.0700;**86.0594** | 30 |
| 12 | β-aminobutyric isolithocholate ester | C<sub>28</sub>H<sub>47</sub>NO<sub>4</sub> | 460.3449 | 3.27 | 4.2 | 341.2844;323.2734;104.0700;**58.0638** | 30 |
| 13 | γ-aminobutyric isolithocholate ester | C<sub>28</sub>H<sub>47</sub>NO<sub>4</sub> | 460.3439 | 1.75 | 4.4 | 104.0700; **58.0638** | 30 |

\*Detected in LCA incubation; <sup>a</sup> Detected at t0; **Bold**: characteristic MS/MS fragment in positive polarity

The re-conjugated secondary BAs were confirmed as (D or L) iso/leucolithocholate ester **(1,2,3)**, D-valolithocholic acid **(4)**, L-valolithocholic acid **(5)**, D-valoisolithocholate ester **(6)**, (D or L)valolithocholate ester **(7)**, L-valoisolithocholate ester **(8)**, aminobutyric lithocholic acid (α, β or γ) **(9)**, 2-Aminobutyric acid isolithocholate ester (α-aminobutyric isolithocholate ester) **(10)**, aminobutyric lithocholic acid **(11)**, 3-Aminobutyric acid isolithocholate ester (β-aminobutytic isolithocholate ester), **(12)**, 4-Aminobutyric acid isolithocholate ester (γ-aminobutyric isolithocholate ester) **(13) (Table 2)**. Four of these MCBAs were identified in fecal samples before the incubation. The leucine conjugates were identified in all volunteers while valine conjugates were only identified in fecal samples of one volunteer before incubations. The aminobutyric acid conjugates were not found in fecal samples before incubations. The iso/leucolithocholic ester **(1,2)** was identified in all volunteers and iso/leucolithocholic ester **(1)** in only one volunteer. In the case of valine conjugates, the D-valoisolithocholate ester **(6)** and valolithocholate ester **(7)** were the MCBAs identified in fecal samples before incubations. Both amides and esters showed lower retention times than the free LCA backbone and therefore a higher polarity. Aminobutyric lithocholic acid (α, β or γ) **(9)** was the MCBA capable of reducing the polarity of LCA the most, from 5.18 to 3.0 min.

Unlike the previous study, the intensity of the MS/MS fragments and the relative abundance were able to discriminate between amidated and esterified MCBAs. The MS/MS fragmentation patterns in both negative and positive polarities were analysed. In the same way, a wide range of collision energies (10-60 ev) were used for screening all fragmentation possibilities as well as a combination of collision energies and fragmentor voltage of the electrospray ionization source. The negative mode was able to confirm in a very powerful and reproducible way the presence of MCBAs because it released the amino acid residue as the major fragment using the range from 20 to 50 ev.9,29,30 The BAs re-conjugated with valine, leucine and aminobutyric acid released a fragment of m/z 116.0717, 130.0873 and 102.0560 as main fragments, respectively **(Fig. 4)**. However, there were no fragments in negative mode able to discriminate amides and esters. The fragmentation study in negative mode also showed characteristic fragments of the MCBAs corresponding to the neutral loss of carbon dioxide (m/z 430.3670 in the case of valine; **Fig. 4)**. The positive polarity results showed characteristic fragments that allowed to confirm the presence of MCBAs. Both amides and esters showed in most of the MCBAs a common fragment with m/z 341.2821 that appears as the characteristic fragment of these molecules in the Competitive Fragmentation Modeling for Metabolite Identification (CFM ID; https://cfmid.wishartlab.com/), and other fragment with m/z 323.2722 corresponding to the loss of a water molecule from the previous fragment. Besides, characteristic fragments corresponding to the protonated amino acids released from the amide and ester molecule were also identified. Only was possible the differentiation of the amides and esters once the released amino acid was fragmented. The results showed that the amides release a characteristic MS/MS fragment related to the water loss (−18) of the amino acid while esters release a fragment related to water plus carbon monoxide loss (−46) in positive polarity. The results showed a characteristic fragment of m/z 100.07 and 72.08 for amides and esters respectively in the case of valine **(Fig. 5)**. The same fragmentation patterns occurred for leucine and aminobutyric isomers **(Fig S1; Fig S2)**. The fragmentation results suggested that the occurrence of the characteristic fragment of the esters corresponding to the water plus carbon monoxide loss is more feasible because the oxygen of the acid group of the amino acid is involved in the ester and somehow is easier to lose it once the amino acid is released. On the other hand, once the amino acid is released from the amide is the amino group the one has to be protonated and is unlikely to remove the complete acid group of the amino acid. This fragmentation behaviour was contrasted with the fragmentation patter of the dipeptides and esters of amino acids by the Competitive Fragmentation Modeling for Metabolite Identification (CFM ID; https://cfmid.wishartlab.com/). Examining the case of the dipeptide composed by valine and leucine (leucyl-valine and valyl-leucine), similar results were found. Depending on the usage of the amino or the carboxylic group for the peptide bond by the amino acids, the main fragment will be the loss of either H2O or H2O+CO respectively. Thus similar as in amidated MCBAs where the amino group of the amino acid is used, the amino acid (valine in the dipeptide leucyl-valine) release the fragments corresponding to the valine and the loss of H2O from valine. This fragmentation pattern was observed as well in the case of valyl-leucine where then the leucine, which is linked through its amino group, loss a molecule of water. Comparing with valine esters, in case of L-valine methyl ester an similarly to esterified MCBAs, when the amino group of the amino acid is free and its acid group is used for the ester bond, the main fragment found is the loss of H2O+CO. Therefore, the MS/MS fragmentation results showed that if the amino group of the amino acids is used for the bond the amino acid will loss a molecule of water, similarly to amidated MCBAs, and in the other hand if the acid group of the amino acid is used for the bond, similarly to esterified MCBAs, the amino acid will lost water plus carbon monoxide.

**Figure 4.**
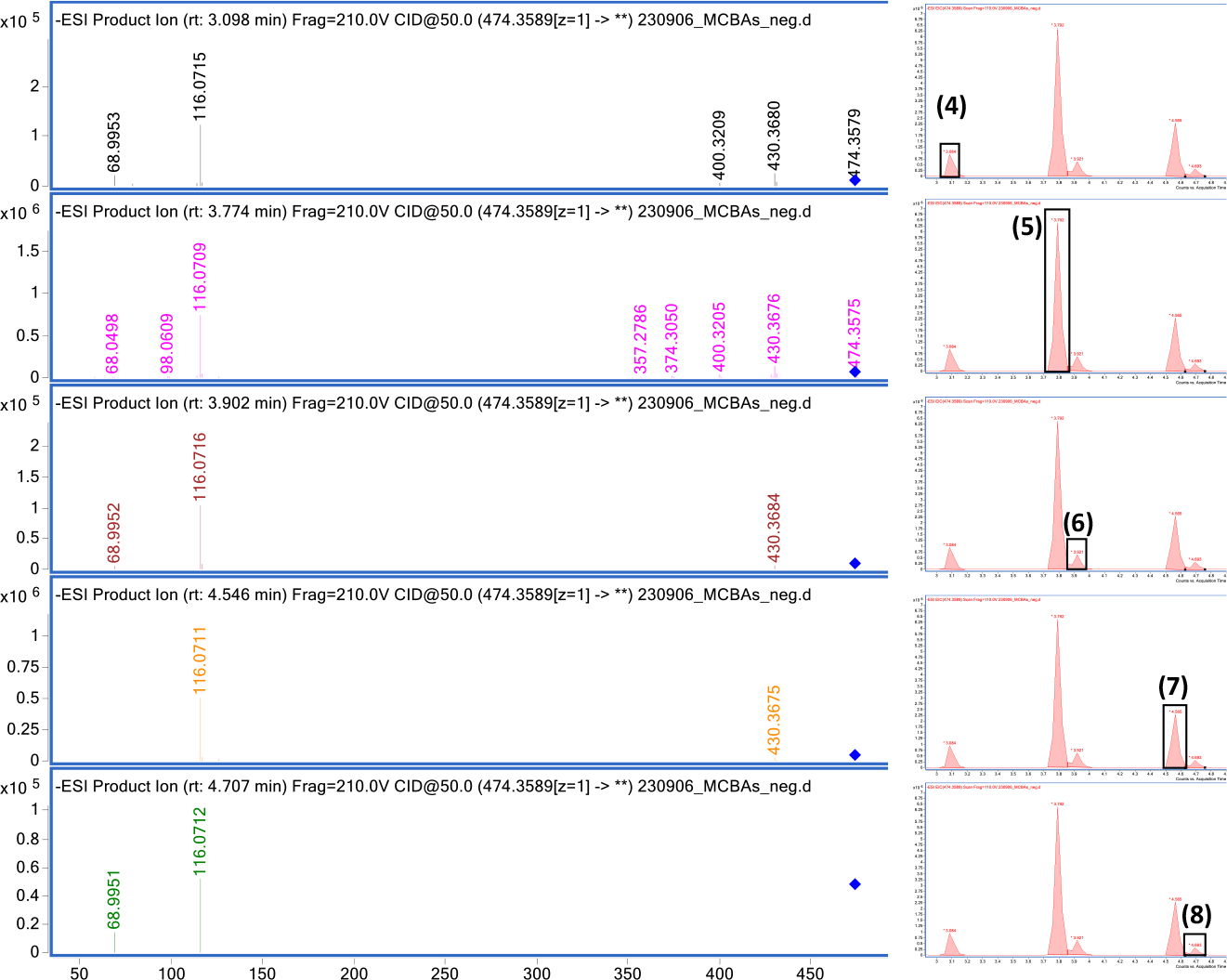
MS/MS spectra fragmentation in negative mode polarity of the five MCBAs valine conjugated. MS/MS spectra corresponding to: (4) D-valolithocholic acid; (5) L-valolithocholic acid; (6) D-valoisolithocholate ester; (7) Valolithocholate ester; (8) L-valoisolithocholate ester.

**Figure 5.**
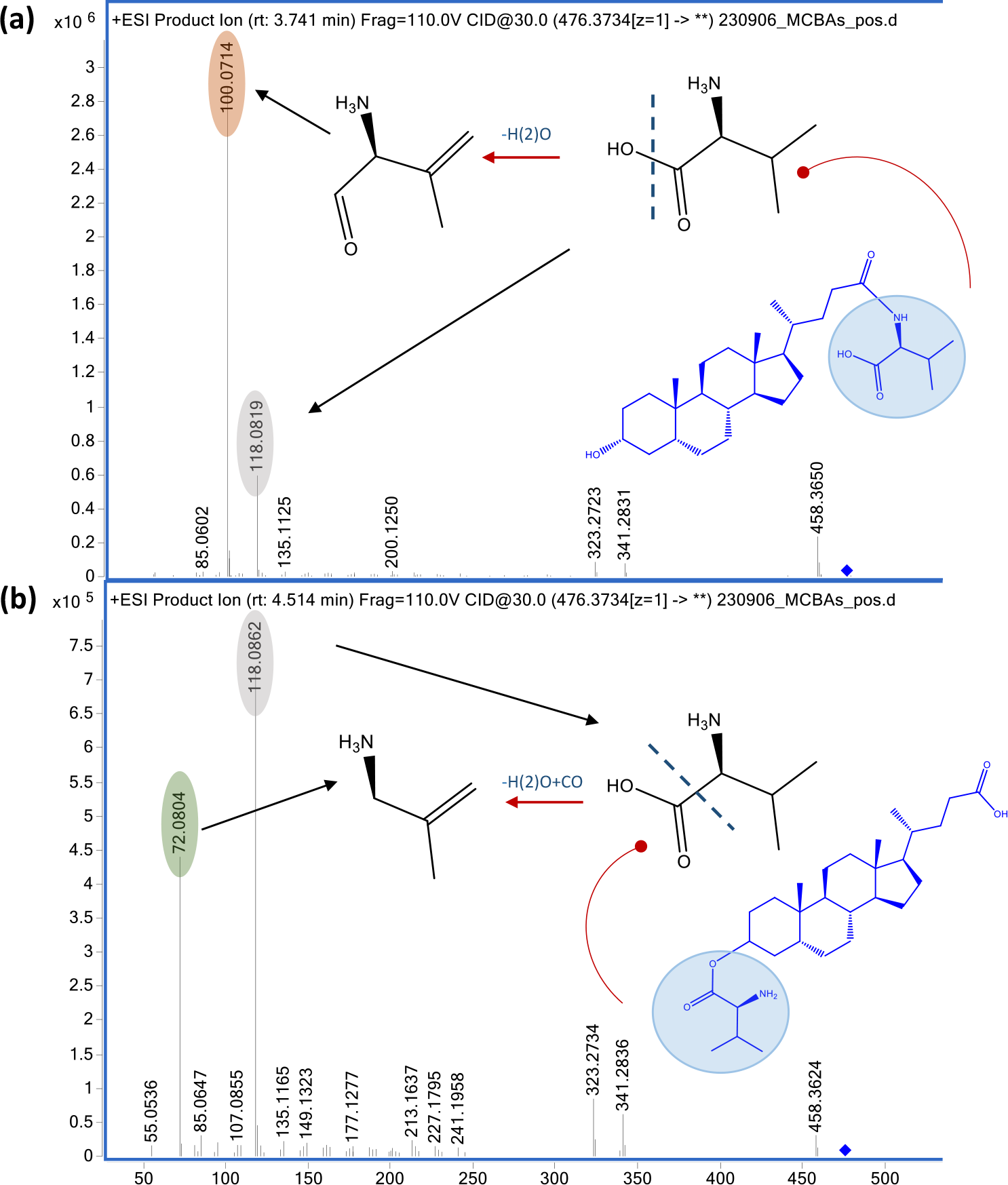
MS/MS spectra fragmentation of the amidated and esterified MCBAs. (a) MS/MS spectra of amides; (b) MS/MS spectra of esters. Amino acid (blue circle); characteristic fragment of the amino acid protonated (grey circle); characteristic fragment of the H(2)O loss related to amides (orange circle); characteristic fragment of H(2)O plus CO loss related to esters (green circle)

The interaction between the gut microbiota and the BAs is the crucial point to understand BAs metabolism and their implications in metabolic disorders. In this study fecal incubation with the primary bile acid, chenodeoxycholic acid, showed the classical conversions to secondary BAs and confirmed the presence of amidated and esterified MCBAs thanks to the fragmentation pattern of the amino acid conjugated. As in previous studies, the primary BAs conversions by the gut microbes lead to produce and accumulate epimers of secondary BAs and LCA. The mechanism to produce β-orientation of the hydroxyl groups confers more hydrophobicity to the molecule and therefore reduces the toxicity.^**31**^ This enzymatic activity is the hydroxysteroid dehydrogenase (HSD) and is performed by bacterial enzymes that act on hydroxyl groups of the BAs. Additionally, the BAs hydrophobicity and therefore toxicity depends on the number of the hydroxyl groups, being LCA the most hydrophobic BA.^**32**^ The results showed the LCA as the major BA backbone substrate to produce the MCBAs. These results suggested a possible specific requirement of the bacteria to avoid the presence of the unconjugated lithocholic acid that has been described as highly toxic.^**33**^ This is similar to what happens with hepatically conjugated BAs. In general, the occurrence of primary BAs conjugated with glycine and taurine confers them specific physiochemical and metabolic properties for preventing their toxicity. These conjugates increase the solubility in acidic pH and being fully ionized at small intestine pH. This prevents the passive absorption of the epithelial cell and passive paracellular absorption by the size and negative charge of the molecule. If BAs were synthetically conjugated with other amino acids such conjugates would be readily hydrolysed by pancreatic carboxypeptidases.^**34**^ Therefore, conjugation with glycine and especially taurine make BAs hardly absorbable. Unconjugated and some glycine-conjugated BAs are reabsorbed via passive diffusion along the small intestine and on the contrary the active transport of BAs occurs in the ileum and passive absorption of hydrophobic secondary BAs occurs in the colon.^**35**^

Additionally, the gut bacteria are able to hydrolyse the conjugated BAs by the bile salt hydrolase activity (BSH), widespread in commensal bacteria colonizing both the small intestine and colon.^**36**^ The BSH activity and therefore, the balance of unconjugated/conjugated BAs may play a key role in improving the control of different metabolic disorders related to cholesterol, lipids and glucose metabolism, bowel disease or cancer. It has been described how the modulation of the BSH activity increases the ileum content of conjugated BAs inhibiting the intestinal FXR-FGF15 signaling pathway and leading to a reduction of the hepatic cholesterol and decreased lipogenesis. The aforementioned relevance of the unconjugated/conjugated BAs and their ability to be absorbed show the importance of the new re-conjugated BAs and the need of an adequate characterization. This study confirmed thirteen MCBAs and could identify and differentiate them for the first time as amides and new esters. The results of the study showed a polarity reduction of the MCBAs compared with the lithocholic acid, especially the aminobutyric lithocholic acid **(9)**. These results suggest that this re-conjugation could prevent the passive absorption of unconjugated and more hydrophobic secondary BAs.

The MCBAs identified were conjugated with branched-chain amino acids (BCAA) and aminobutyric acids derivatives. The function of these re-conjugated BAs needs to be revealed and therefore the relationship with the amino acid conjugated. The link between the gut microbiota and the abundance of the BCAA levels and insulin resistance have been studied showing an increase of BCAA in studies where the obesity and diabetes model were evaluated.^**37**^ We hypothesized that non-healthy individuals, because of the gut dysbiosis, are not able to produce the re-conjugated derivatives with the BCAA and show an increase of free BCAA in the fecal metabolome. Regarding aminobutyric acid derivatives, some strains of bacteria have demonstrated the ability to produce gamma aminobutyric acid (GABA) from glutamate in the human intestinal tract.^**38,39**^ Therefore, the re-conjugation of BAs with aminobutyric acid derivatives may be possible.

## Conclusion

The relevance of the BAs in lipid and glucose metabolism supports the need to develop suitable methods to evaluate them. In particular, the re-conjugation of BAs by the gut microbiota, little studied so far, is an important step for a full knowledge of the BAs profile. Besides, the differentiation between MCBAs with esters and amide bonds could open new opportunities to evaluate the impact of the BAs metabolism on health.

## Author Contributions

Carlos J. Garcia: conceptualization, formal analysis, investigation, methodology, software, validation, writing – original draft, writing – review & editing. Rocio Garcia Villalba: validation, supervision, writing – review & editing. David Beltrán: investigation, methodology; Maria D. Frutos-Lisón. Francisco A. Tomás-Barberán: Funding acquisition, supervision, writing – review & editing.

