## Supplementary material for "MS/MS fragmentation pattern analysis confirms the production of the new esterified bile acids by the human gut microbiota": Fig S1

**Supplementary Figure 1**

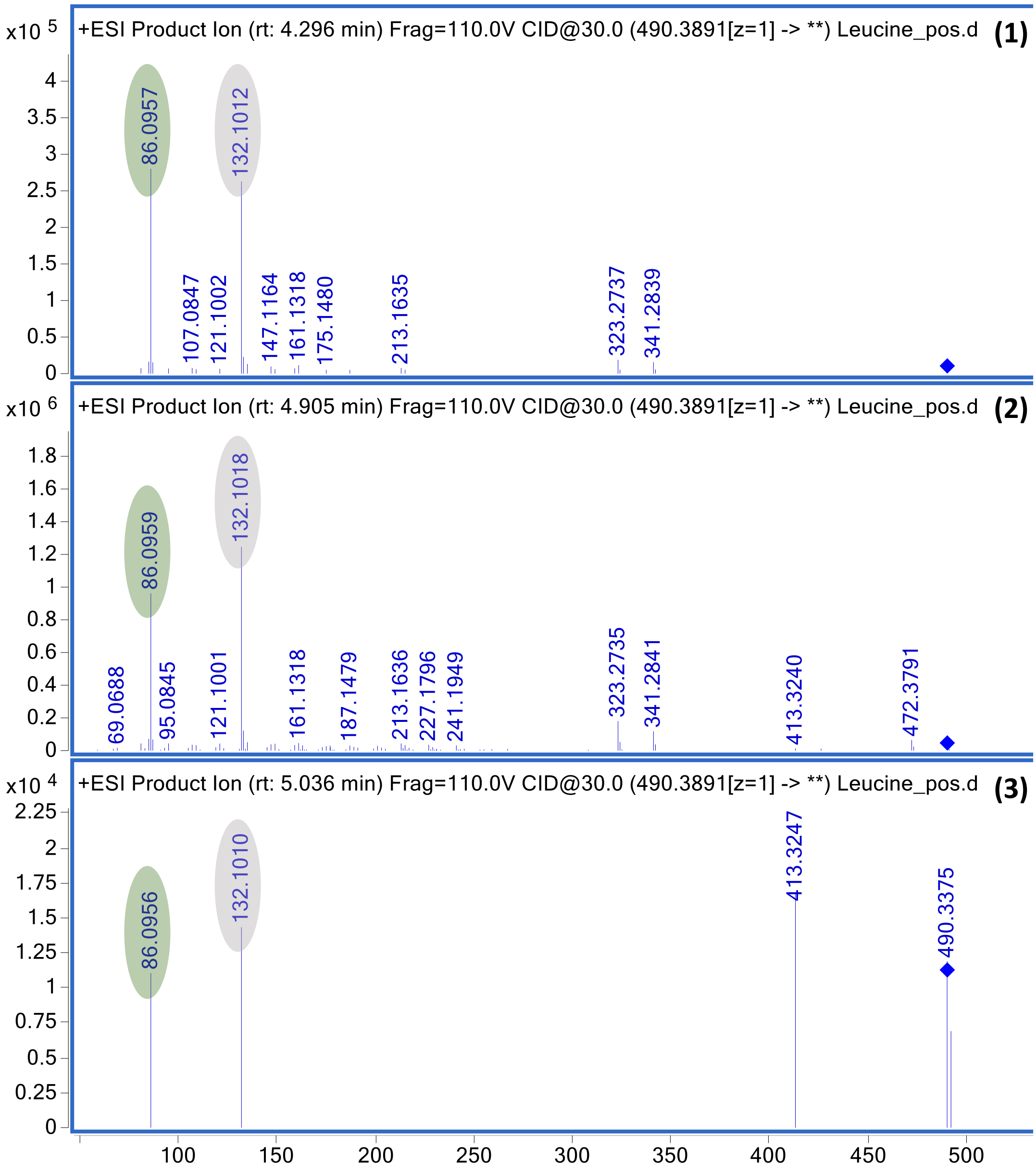


**Supplementary Figure 1.** MS/MS spectra of amidates and esterified MCBAs with leucine. Esters (blue MS/MS spectra); Characteristic fragment of the amino acid protonated (grey circle), characteristic fragment of H(2)O plus CO loss related to esters (Green circle); (1) iso/leucine(L or D)lithocholate ester; (2) ) iso/leucine(L or D)lithocholate ester; (3) iso/leucine(L or D)lithocholate ester.
