## Supplementary material for "MS/MS fragmentation pattern analysis confirms the production of the new esterified bile acids by the human gut microbiota": Fig S2

### Supplementary Figure 2

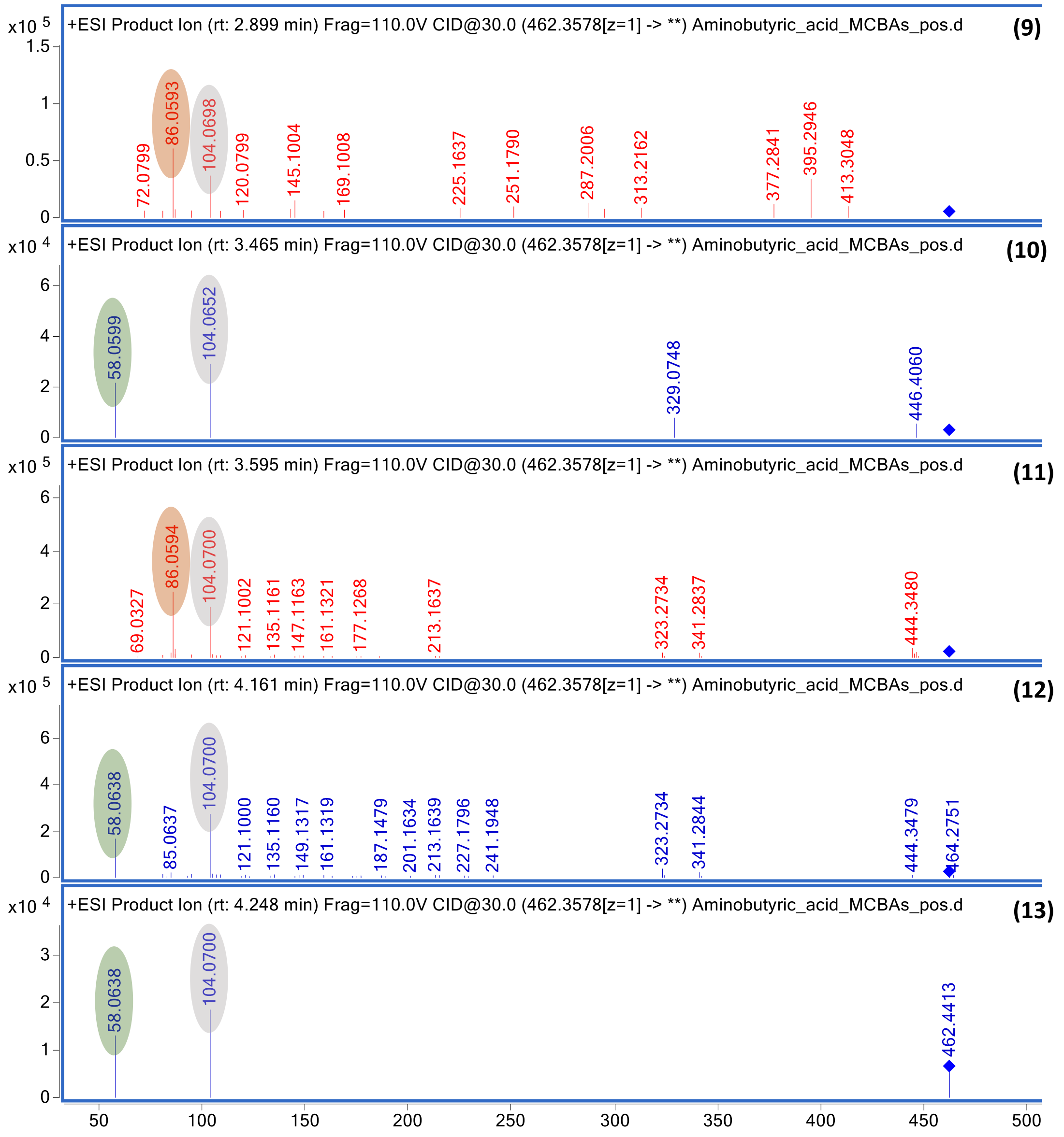


**Supplementary Figure 2.** MS/MS spectra of amidates and esterified MCBAs with aminobutyric acid. Amides (red MS/MS spectra), esters (blue MS/MS spectra). Characteristic fragment of the amino acid protonated (grey circle); characteristic fragment of the H(2)O loss related to amides (orange circle), characteristic fragment of H(2)O plus CO loss related to esters (Green circle). (9) Aminobutyric lithocholic acid; (10) α-aminobutyruc isolithocholate ester; (11) Aminobutyric lithocholic acid; (12) β-aminobutytic isolithocholate ester; (13) γ-aminobutyric isolithocholate ester
